## Supplementary Material for "Molecular mechanism of Afadin substrate recruitment to the receptor phosphatase PTPRK via its pseudophosphatase domain"

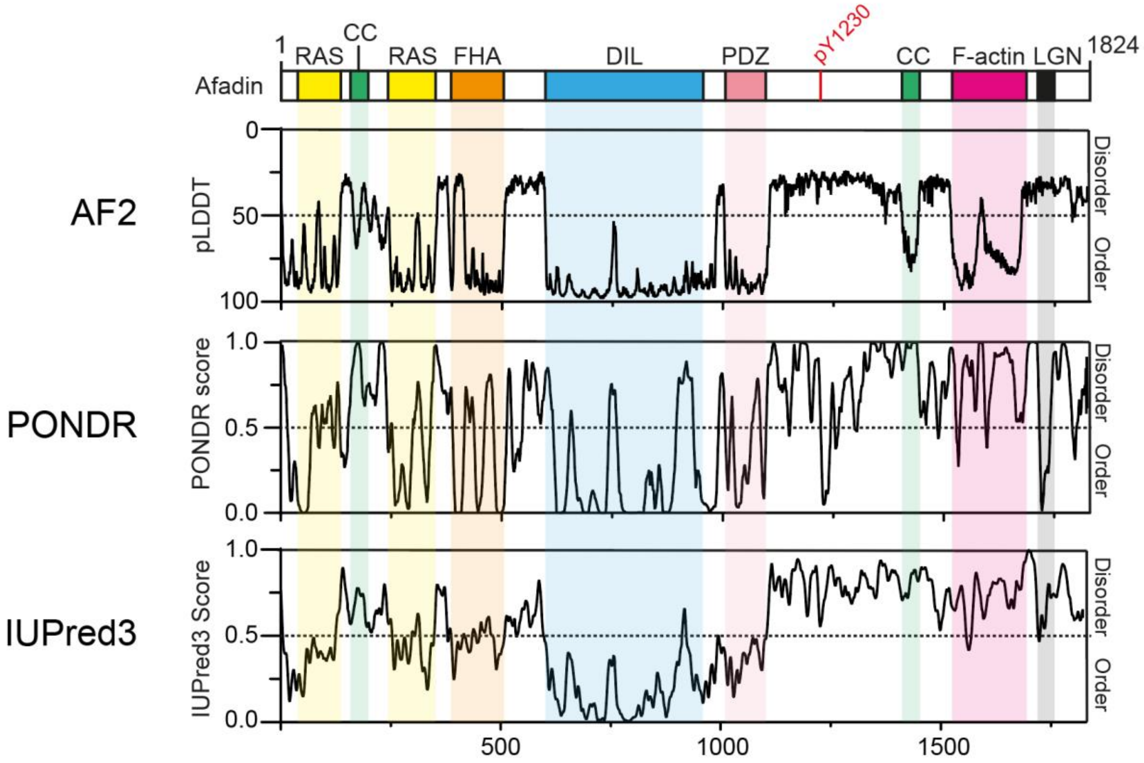


**Supplementary Figure 1: Disorder predictions for Afadin.** Top: Schematic of full-length human Afadin (also known as AF6 or MLLT4), with domain annotation based on UniProt ID: P55196, colored as in main Figure 1A. The predicted local distance difference test (pLDDT) for the Afadin AlphaFold2 (AF2) prediction from the AF Protein Structure Database (ID: P55196, retrieved 7/02/2022) is aligned to disorder predictions from the PONDR and IUPred3 servers, highlighting the agreement of pLDDT scores with dedicated disorder prediction methods. For clarity, the Y axis for the AF2 pLDDT plot has been inverted, so as to have predicted disorder consistently displayed in the top half of all graphs.

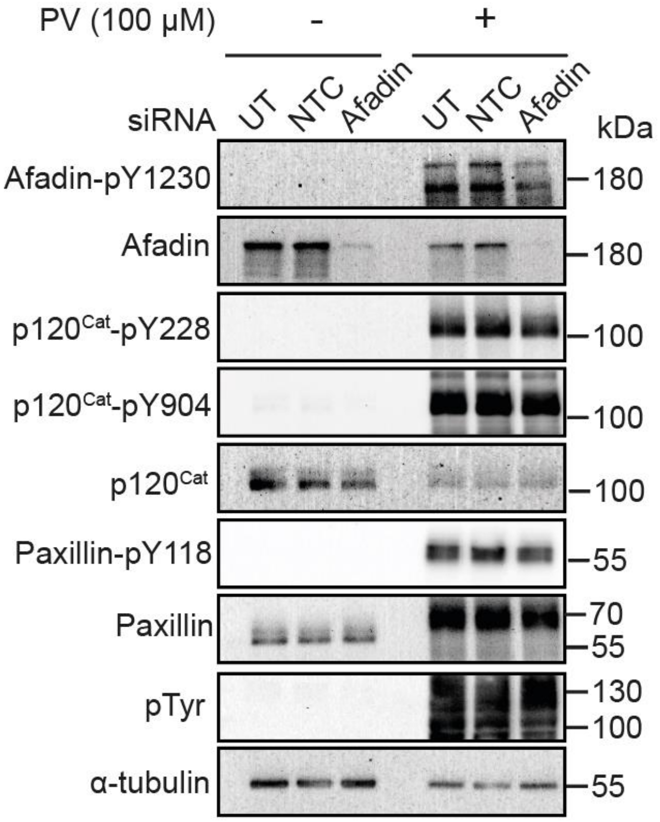


**Supplementary Figure 2: Afadin-pY1230 antibody validation.** Immunoblot analysis of MCF10A cells either untransfected (UT) or transfected with non-targeting control (NTC) and *AFDN* (Afadin) targeting small-interfering (si)RNAs. Phosphorylation of cellular proteins was stimulated by treatment with 100 µM sodium pervanadate (PV) for 30 min prior to cell lysis.

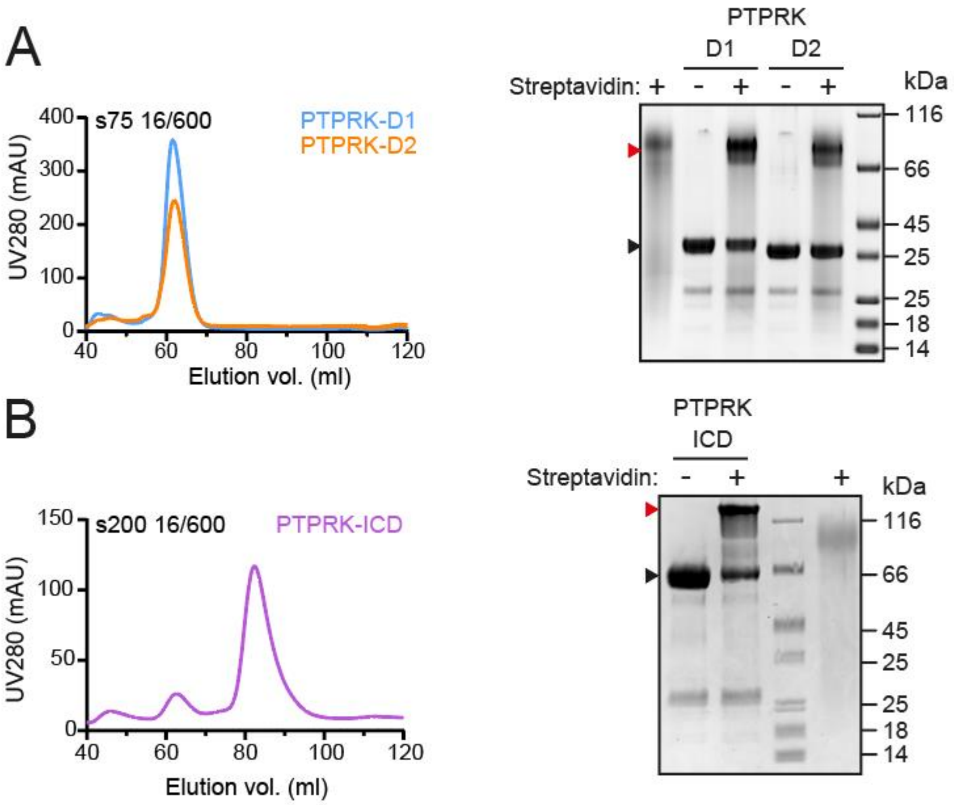


**Supplementary Figure 3: Purification of *in vivo* biotinylated PTP domains.** (**A**) Left: Following Ni-NTA affinity chromatography, *in vivo* biotinylated PTPRK D1 and D2 domains were purified by size-exclusion chromatography (SEC) on a S75 16/600 column. Right: After SEC purification, *in vivo* biotinylated PTPRK D1 and D2 domains were incubated with or without streptavidin, resolved by SDS-PAGE and visualized by Coomassie staining. The mobility shift upon streptavidin binding to biotinylated protein is indicated by a red arrowhead. (**B**) Left: Following Ni-NTA affinity chromatography, *in vivo* biotinylated PTPRK-ICD was purified by SEC on a S200 16/600 column. Right: After SEC purification, biotinylation of PTPRK-ICD was assessed as described in (A).

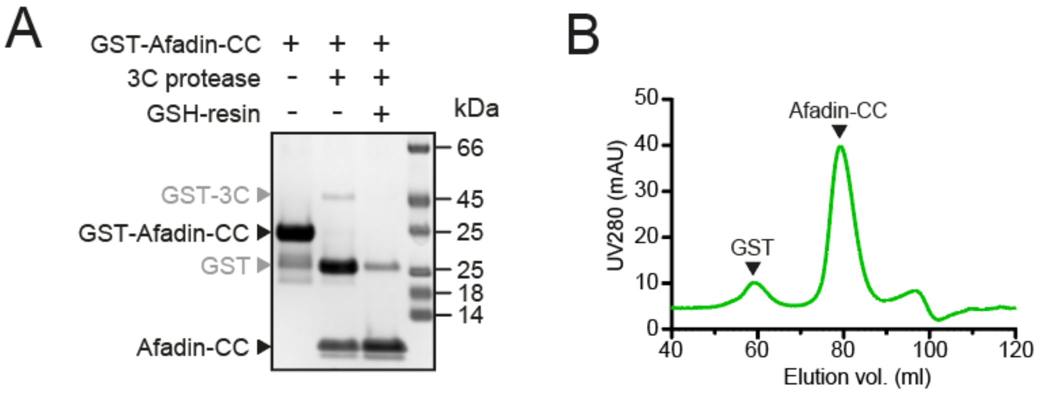


**Supplementary Figure 4: GST-tag removal from Afadin-CC for SEC-MALS and ITC experiments.** (**A**) GST was removed from GST-Afadin-CC by overnight cleavage with GST-3C protease. Cleaved GST and GST-3C were removed by incubation with GSH-sepharose 4B (GSH-resin). (**B**) Residual cleaved GST was removed by SEC of the Afadin-CC sample on a S75 16/600 column.

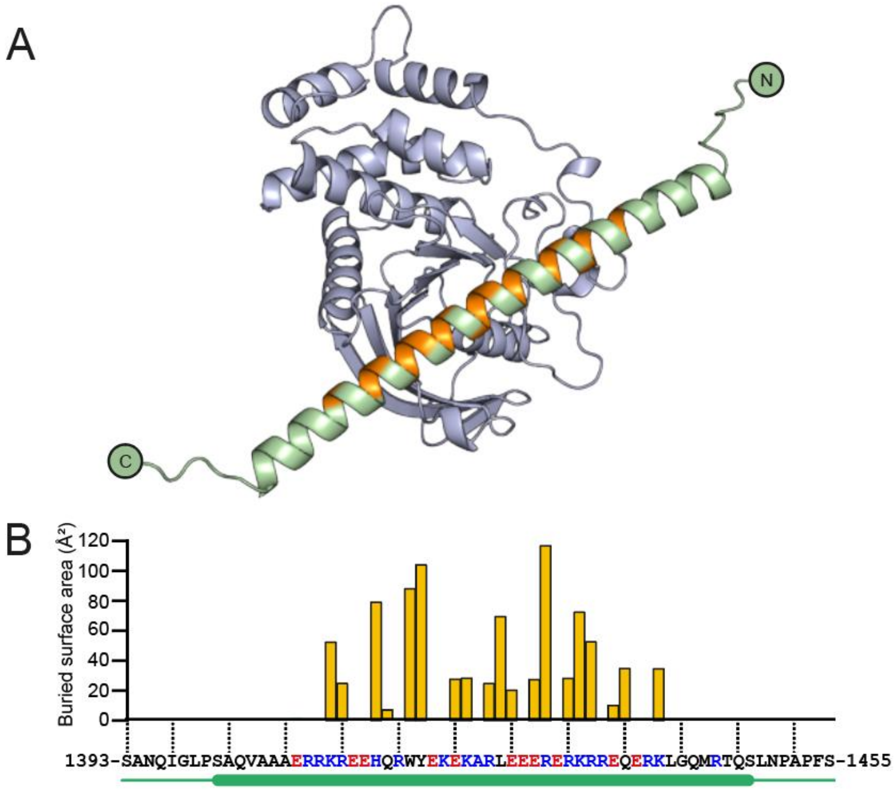


**Supplementary Figure 5: Structural prediction of the PTPRK-D2:Afadin-CC complex.** (**A**) Ribbon diagram of the AF2-Multimer prediction for the PTPRK-D2 domain (blue) in complex with the full Afadin-CC (aa. 1393-1455, green). Afadin-CC residues that contribute to the interaction interface are highlighted in orange. (**B**) Graph of buried surface area (Å$^2$) per residue of Afadin-CC for the PTPRK-D2:Afadin-CC complex. All residues which contribute to the interface are within the core sequence of charged residues (highlighted red/blue for acidic/basic, respectively) within the helical region of Afadin-CC (highlighted by green cylinder). Buried surface area calculations were perfomed using the 'Protein interfaces, surfaces and assemblies' service (PISA) at the European Bioinformatics Institute (Krissinel and Henrick, 2007).

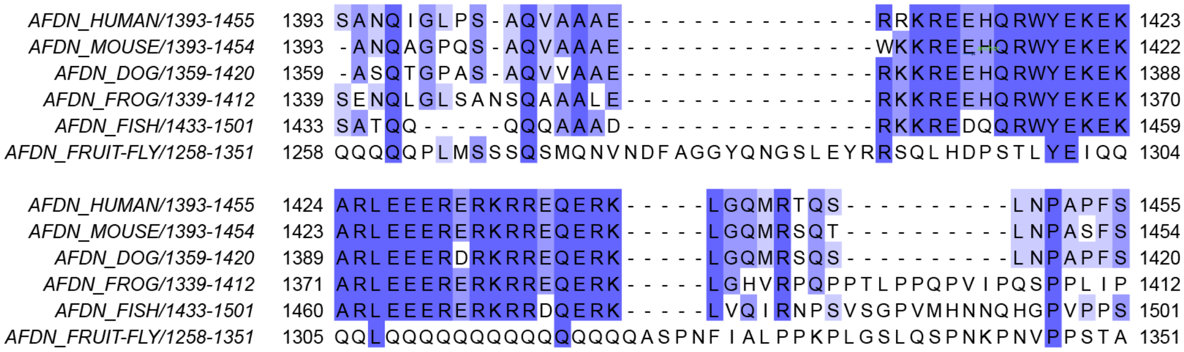


**Supplementary Figure 6: Species conservation of Afadin-CC.** Multiple sequence alignment of the equivalent Afadin-CC region (hsAFDN aa. 1393-1455) from human (*Homo sapiens*) mouse (*Mus musculus*), dog (*Canis familiaris*), frog (*Xenopus laevis*), zebrafish (*Danio rerio*) and fruit fly (*Drosophila melanogaster*).

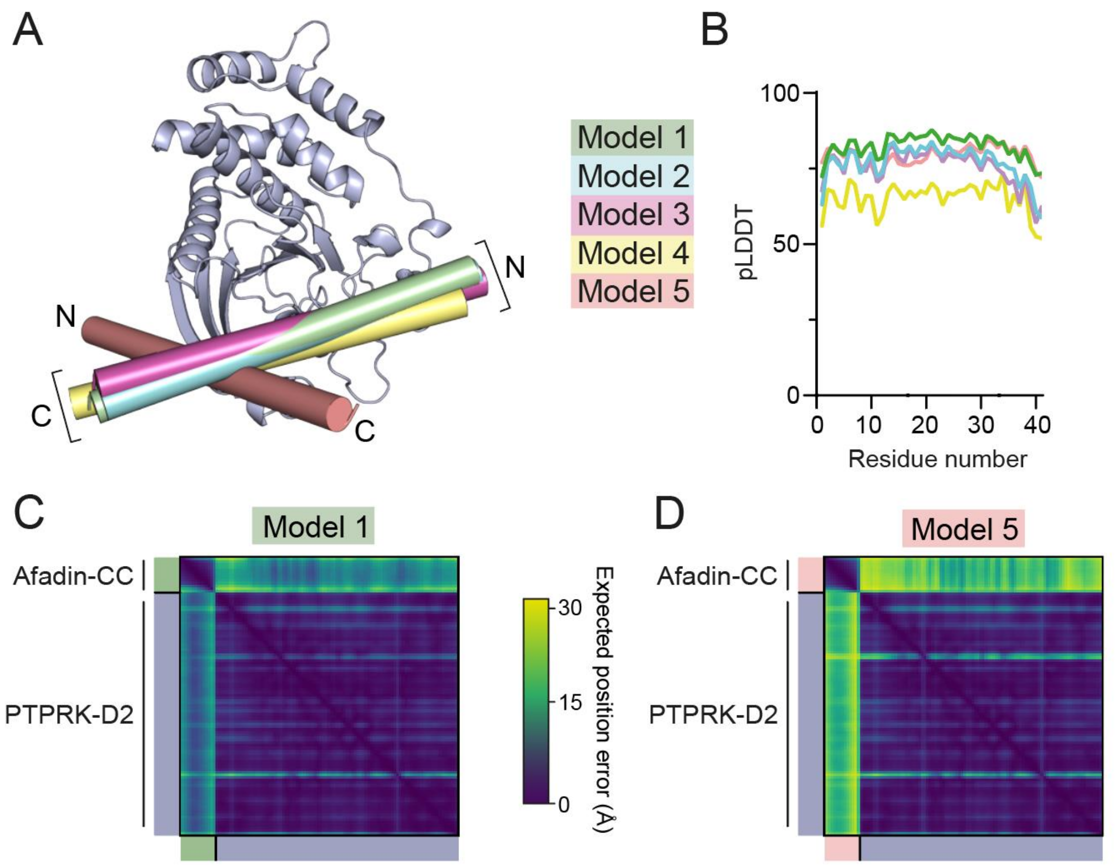


**Supplementary Figure 7: Prediction quality analysis of AF2 multimer generated PTPRK-D2:Afadin-CC complex models. (A)** The 5 AF2-Multimer generated PTPRK-D2:Afadin-CC models were superposed using the PTPRK-D2 chain only. The Afadin-CC chains of each ranked model are shown as cylinders (colored as indicated) with N- and C-termini highlighted. **(B)** Plot of predicted local distance difference test (pLDDT) for the Afadin-CC chain of each ranked PTPRK-D2:Afadin-CC model, colored as in (A). **(C-D)** Predicted aligned error (PAE) plot for highest (**C,** model 1) and lowest (**D**, model 5) ranked models for the PTPRK-D2:Afadin-CC complex. The high expected position error for Afadin-CC vs PTPRK-D2 residues in Model 5 indicates a low confidence in the relative positioning of these domains for the lower ranked models vs that observed for Model 1.

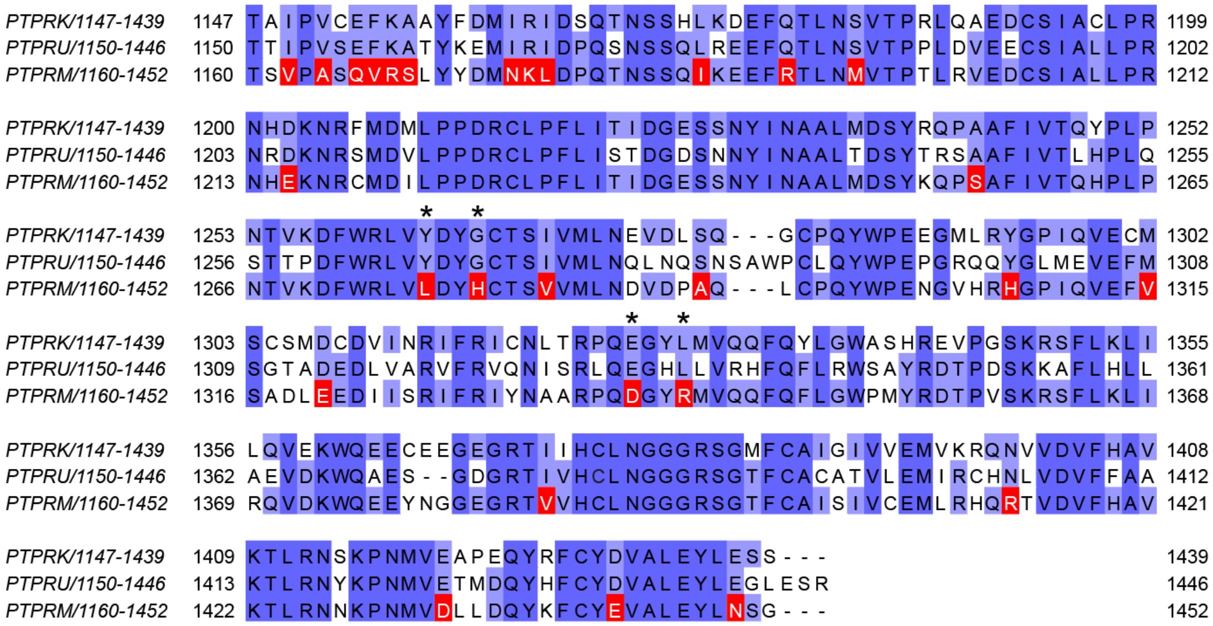


**Supplementary Figure 8: Mapping of unique PTPRM residues.** Multiple sequence alignment of human PTPRK, PTPRU and PTPRM D2 domain sequences. Residues which are conserved in both PTPRK and PTPRU, but differ in PTPRM, which were used for interaction site mapping, are highlighted in red. Asterisks denote divergent PTPRM residues present at the PTPRK-D2:Afadin-CC interface, as illustrated in main Figure 3D and E.

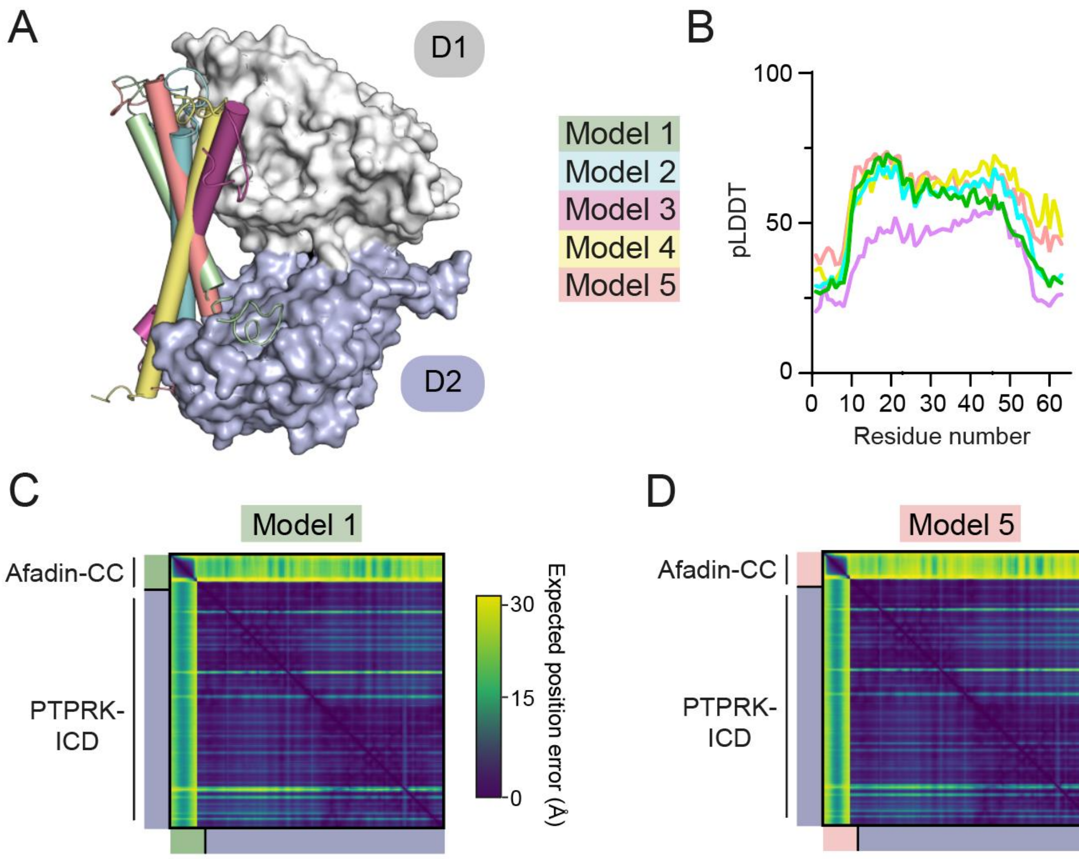


**Supplementary Figure 9: Prediction quality of models prior to biochemical mapping (A)** The 5 AF2-Multimer generated PTPRK-ICD:Afadin-CC models were superposed using the PTPRK-ICD chain only. The Afadin-CC chains of each ranked model are shown as cylinders (colored as indicated) **(B)** Plot of predicted local distance difference test (pLDDT) for the Afadin-CC chain of each ranked PTPRK-ICD:Afadin-CC model, colored as in (A). These scores are markedly lower than for the model generated using the PTPRK-D2 (see Supplementary Figure 6) **(C-D)** Predicted aligned error (PAE) plot for highest (**C,** model 1) and lowest (**D**, model 5) ranked models for the PTPRK-D2:Afadin-CC complex. Both top and bottom ranked models generated using the PTPRK-ICD have worse PAE scores when compared to the PTPRK-D2 models (see Supplementary Figure 6).

### Supplementary References

Krissinel, E., and Henrick, K. (2007). Inference of macromolecular assemblies from crystalline state. J Mol Biol *372*, 774-797.
